# Causal modeling refutes Hockett’s hypothesis that bite configuration affects human sound evolution

**DOI:** 10.1101/2020.02.20.957407

**Authors:** Sergei Tarasov, Josef C. Uyeda

## Abstract

Blasi *et al.* (Research article, 15 March 2019) argue in support of Hockett’s hypothesis – that languages of hunter-gatherers are less likely to develop labiodental sounds than those of agricultural societies due to the former’s heavy-wear diet that favors edge-to-edge bite, thereby decreasing likelihood of labiodental articulation. We reanalyze the data in Blasi *et al.* and find little to no support in favor of the Hockett’s hypothesis. The negative association between labiodentals and hunter-gatherers instead appears to be an artifact of labiodental decline with increasing distance from Africa, which is a general trend of language phonemes.

---

In their world-wide analysis of human languages, Blasi *et al.* (*1*) conclude that subsistence type is (i) a significant predictor of the labiodental counts using regression models that correct for language family, area and number of nonlabiodental phonemes, and that (ii) languages of hunter-gatherers appear to have substantially fewer labiodentals than those of agrarian food-producers. While regression models are powerful statistical tools for inferring correlation between variables, it is of course true that correlation does not mean causation. The distribution of hunter-gatherers is shaped by environmental (*2*) and historical factors (*3*) and is therefore non-uniform across geographic areas. Similarly, languages of varying families might have inherited different potential to evolve labiodentals due to their ancestry that was driven by historical human dispersal and hence, also associated with geography. Thus, the correlation between labiodentals and subsistence might be explained by their mutual dependence on geography rather than the effect of diet on the ability to produce labiodentals. Regression analysis alone cannot disentangle these different types of dependencies, and can often result in overstating the evidence for a causal relationship.

We reanalyzed both datasets of Blasi *et al.* – GMR and AUTOTYP – by adopting Bayesian inference (*4*) for two causal models, in which the presence of labiodentals was set as either an independent or dependent binomial proportion with respect to subsistence type (Fig. 1A). In both models, observations were conditioned on geographic area to control for its effect. If Hockett’s hypothesis is true, then the dependent model, indicating causality between subsistence and labiodentals, is expected to be selected across all regions. We used Bayes factors (BFs) to assess model fit and found strong support for the dependent model only in North-Central Asia [BF: GMR = −5.6, AUTOTYP = −2.0], and weak support in Africa [BF: GMR = −1.5, AUTOTYP = 0.6]. In the remaining eight areas, the independent model exhibits similar or better fit [-0.004 *≤* BFs *≤* 2.4; Fig. 1A], thus contradicting to Hockett’s hypothesis.

**Figure 1:**
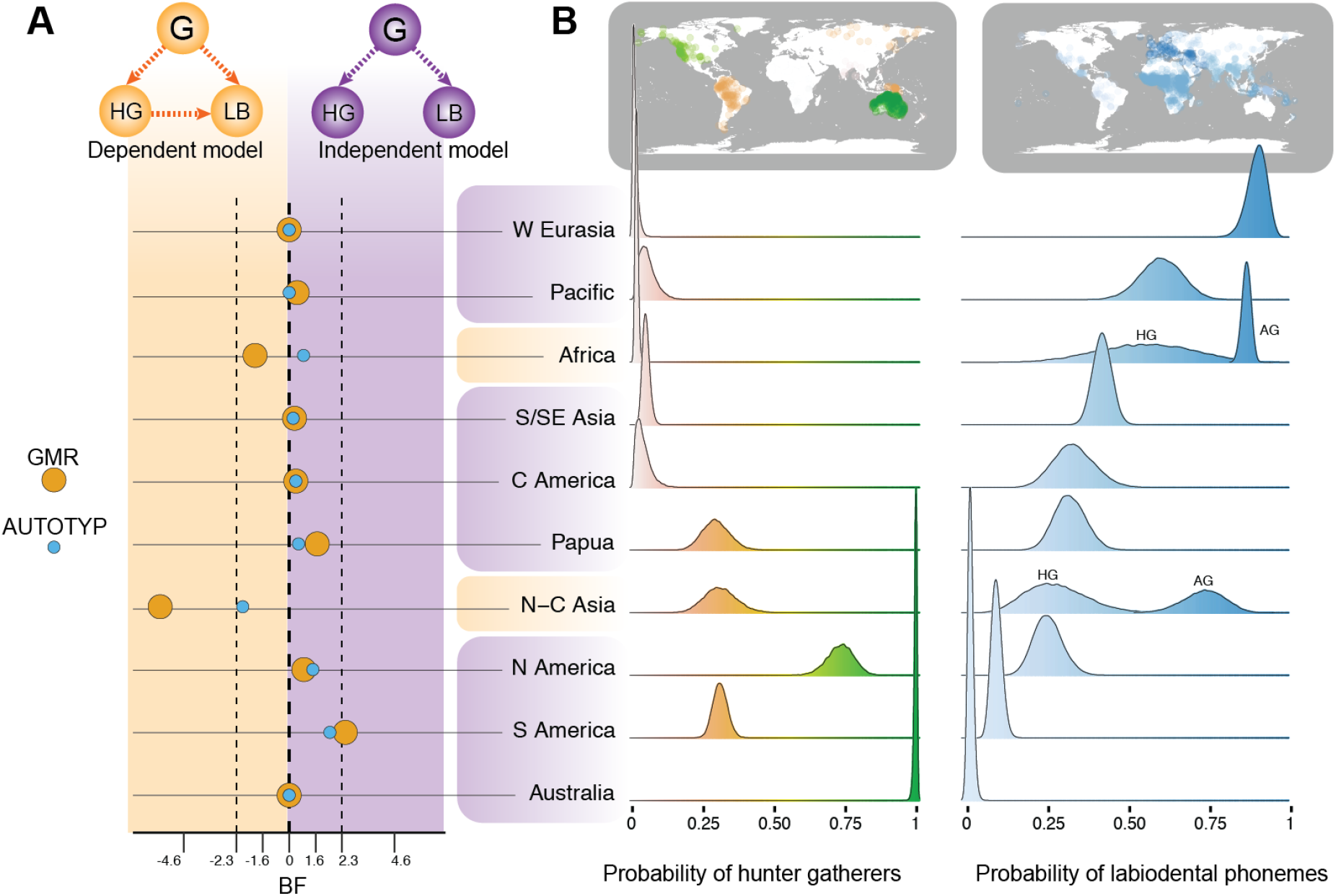
Distribution of hunter gatherers and labiodentals across geographic regions. **(A)** Statistical fit of dependent (top, left) and independent (top, left) causal models shown using logarithm of Bayes factor *BF* for GMR and AUTOTYP datasets; in the dependent model the proportion of labiodentals (LB) is affected by both geography (G) and hunter gatherers (HG), while in the independent model it is affected only by geography; Africa and North-Central Asia are the only regions that support the dependent model. **(B)** Posterior distribution for the probabilities of hunter gatherers and labiodentals inferred using best-fit causal models (density plots). Africa and North-Central Asia have bimodal distributions of labiodentals since they support the dependent model in which labiodentals are modeled separately for hunter gatherers (HG) and agricultural societies (AG).

We do, however, find that the proportions of languages with labiodentals and hunter-gatherer subsistence are globally correlated in our worldwide analysis [BF: GMR = −213.2, AUTOTYP = −18.1] and vary inversely among regions (rather than within regions) given linear regression [GMR dataset: slope = −0.6, 95% credible interval (−1.3, 0.0); AUTOTYP dataset: slope = - 0.5, 95% credible interval (−1.3, 0.3); Fig. 1B]. Does this inverse relationship corroborate the evidence of causal relationships between labiodentals and subsistence? Or could such a relationship represent an artifact of the confounding effect of geography? To test this, we ran posterior predictive simulations using the original predictors and an additional one – distance from Africa. The latter was chosen because phonemic diversity declines with a distance from Africa due to serial founder effects (*5*) that may also affect labiodental counts. Our simulations reveal that distance alone accurately explains the observed across-region variation of labiodentals. The predictive power, measured as an average mean squared error, was two-times better for distance (GMR = 0.4, AUTOTYP = 0.8) as it was for subsistence (GMR = 0.9, AUTOTYP = 1.6).

Thus, we argue that the world-wide association between labiodentals and hunter gatherers (except for within Africa and North-Central Asia) reported by Blasi *et al.* can primarily be attributed to across-area variation in labiodental frequency, largely consistent with serial founder effects. The source of dependence of labiodentals on subsistence within Africa and North-Central Asia may indeed be related to Hockett’s hypothesis, but the inconsistent signals in other regions provide evidence that such a conclusion is premature. Rather, we suggest that the dependence of labiodentals in Africa and North-Central Asia can be induced by the confounding effects of phylogenetic non-independence of languages and historical human dispersal. Further research, employing fully phylogenetic framework to language evolution is needed to disentangle the source of dependency in these two regions.

As another evidence for Hockett’s hypothesis, Blasi *et al.* reconstructed ancestral character states for ten labiodental sounds on the phylogenies of Indo-European languages and found that the probability of acquiring labiodental states gradually increased from root to tips. This increase, started 6000-3500 years ago, is argued to have resulted from the spread of softer diet during post-Neolithic. The models, used for ancestral reconstruction in Blasi *et al.*, were traditional continuous-time Markov chains that did not include any data on diet itself, which would enable an explicit test as to whether diet is correlated with changes in labiodental sounds over time (*6*). We find it is unconvincing to state that reconstructing low probability of labiodentals at root should indicate anything about its correlation with diet. Instead, it may mean that the ancestor of Indo-European languages inherited no or few labiodentals from its parent, while the successive increase of probability resulted from the stochastic nature of Markov processes that converge to equilibrium as time elapses.

We propose, if an increase in labiodental sounds was driven by changes in diet, then the latter should have a measurable effect on the labiodental evolutionary rate that should vary across branches of phylogeny to be associated with changes in subsistence. To test this hypothesis, we re-analyzed the original datasets using Bayesian approach (*7*) and adopting stochastic models that allow branch-specific evolutionary rates (*8*). Both datasets yielded no rate shifts or heterogeneity (with posterior probability = 1.0), suggesting that there is little to no evidence that the origin of soft diet corresponded with an increase of labiodental counts in Indo-European languages.

Blasi *et al.* ground their conclusion on evidence from ethnography, historical linguistics and speech biomechanics. We demonstrated that the first two sources do not support their findings and Hockett’s hypothesis. While the biomechanics seems to be a plausible mechanism that may ultimately explain labiodental evolution, our re-analyses suggest that other factors that covary with geography and phylogeny likely play a larger role in driving global patterns of sound changes in human.

## Acknowledgments

We acknowledge constructive communications with Damian Blasi.

## Data and materials availability

All data needed to assess the conclusions of the paper are present in the paper. Scripts and description of analyses related to this paper may be found at https://github.com/sergeitarasov/Tarasov_Uyeda_SupplementaryMaterials_2019.

